# Studying the cellular basis of small bowel enteropathy using high-parameter flow cytometry in mouse models of primary antibody deficiency

**DOI:** 10.1101/2024.01.25.577009

**Authors:** Ahmed D. Mohammed, Ryan A.W. Ball, Amy Jolly, Prakash Nagarkatti, Mitzi Nagarkatti, Jason L. Kubinak

## Abstract

**Background:** Primary immunodeficiencies are heritable defects in immune system function. Antibody deficiency is the most common form of primary immunodeficiency in humans, can be caused by abnormalities in both the development and activation of B cells, and may result from B-cell-intrinsic defects or defective responses by other cells relevant to humoral immunity. Inflammatory gastrointestinal complications are commonly observed in antibody-deficient patients, but the underlying immune mechanisms driving this are largely undefined.

**Methods:** In this study, several mouse strains reflecting a spectrum of primary antibody deficiency (IgA^-/-^, Aicda^-/-^, CD19^-/-^ and J_H_^-/-^) were used to generate a functional small-bowel-specific cellular atlas using a novel high-parameter flow cytometry approach that allows for the enumeration of 59 unique cell subsets. Using this cellular atlas, we generated a direct and quantifiable estimate of immune dysregulation. This estimate was then used to identify specific immune factors most predictive of the severity of inflammatory disease of the small bowel (small bowel enteropathy).

**Results:** Results from our experiments indicate that the severity of primary antibody deficiency positively correlates with the degree of immune dysregulation that can be expected to develop in an individual. In the SI of mice, immune dysregulation is primarily explained by defective homeostatic responses in T cell and invariant natural killer-like T (iNKT) cell subsets. These defects are strongly correlated with abnormalities in the balance between protein (MHCII-mediated) versus lipid (CD1d-mediated) antigen presentation by intestinal epithelial cells (IECs) and intestinal stem cells (ISCs), respectively.

**Conclusions:** Multivariate statistical approaches can be used to obtain quantifiable estimates of immune dysregulation based on high-parameter flow cytometry readouts of immune function. Using one such estimate, we reveal a previously unrecognized tradeoff between iNKT cell activation and type 1 immunity that underlies disease in the small bowel. The balance between protein/lipid antigen presentation by ISCs may play a crucial role in regulating this balance and thereby suppressing inflammatory disease in the small bowel.

## Introduction

There are over 300 clinically recognized forms of primary immunodeficiency, which are heritable disorders of immune system function. The most common manifestation is a deficiency in the production of antibodies (primary antibody deficiencies (PADs)^1,2^. Over two dozen susceptibility genes have been identified that may contribute to primary antibody deficiency, with most being associated with defects in B cell receptor (BCR) signaling or co-stimulatory pathways^1-4^. The degree of antibody deficiency covaries with both the incidence and severity of potential clinical complications. Specifically, gene defects impacting a single antibody class tend to result in less clinically severe phenotypes, while those negatively impacting two or more antibody classes result in the most clinically-severe complications^4,1^. For example, selective IgA deficiency (IgA-deficient only) is generally considered a clinically benign form of antibody deficiency, whereas common variable immunodeficiency (CVID)(deficient in IgA and IgG, and sometimes IgM) results in higher incidence and severity of clinical complications.

Primary immunodeficiency results in immune dysregulation, which is broadly defined as an abnormal immune phenotype. However, this definition is inadequate for three reasons. First, it is an ambiguous phenotype that provides little information regarding underlying defect(s) and no information on their pleiotropic effects. Immunity is an emergent property of dynamic interactions between cells and our understanding of it requires the development of diagnostic techniques that better capture this complexity. Second, it is generally assumed that the degree of immune dysregulation positively correlates with the incidence and/or severity of clinical complication. Empirical support for this assumption is lacking. Third, if immune dysregulation is predictive of disease incidence and/or severity then it would be a valuable biomarker to assess risk in patients. Development of a formal method to comprehensively estimate immune dysregulation would fill these gaps in our knowledge.

The mucosa of the gastrointestinal tract serves as a primary physical and immunological barrier to environmental antigens. Because of the tremendous antigenic load in the gut, immune cells are found in their highest abundance in these tissues^5,6^. Enteropathies are commonly observed in antibody-deficient patients^7,8^, with up to 50% developing clinically significant gastrointestinal symptoms^9^. While much attention has been given to understanding mechanisms of inflammatory disease of the colon, mechanisms of inflammatory pathogenesis in the small bowel (hereafter termed ‘small bowel enteropathy’) are poorly understood. This is a major gap in our knowledge as several inflammatory diseases with small-bowel-specific involvement, including crohn’s disease (CD)^10^ and celiac disease^11^, have been associated with defects in humoral immunity^8,12,13^. Notably, CVID may also be associated with small-bowel-specific involvement termed “CVID enteropathy”^9,14-16^.

Understanding the cellular basis of inflammatory gastrointestinal disease in humans has been hampered by the difficulties in obtaining sufficient materials to study from patients. Thus, it is imperative that assays be developed that maximize the amount of information that can be gleaned from them. Recent advances in flow cytometry, specifically the development of spectrally-equipped flow cytometers, have dramatically increased the number of cellular parameters that can be simultaneously detected in a single tissue sample allowing investigators to generate high-dimensional datasets of functional immune readouts in a tissue-specific manner.

Here, we sought to demonstrate the utility of this technology by exploring the cellular drivers of small bowel enteropathy in mouse models of antibody deficiency. To do this, we developed a small-bowel-specific high-parameter flow cytometry assay incorporating the use of spectrally- and non-spectrally-equipped BD FACSymphony A5 cell analyzers available to us at the University of South Carolina School of Medicine. Using this assay, we describe the initial attempt by our group to construct a small-bowel-specific cellular atlas of immune cell, intestinal epithelial cell (IEC), and intestinal stem cell (ISC) phenotypes. Using this small-bowel-specific cellular atlas, we then compare variation in immune responses and IEC/ISC phenotypes across a series of antibody-deficient mouse models. From these analyses, we outline a novel strategy for quantifying and characterizing immune dysregulation among immunodeficient individuals and highlight a potentially novel axis of disease that may contribute to the pathogenesis of small bowel enteropathy.

## Materials and Methods

### Mouse models

A long-term breeding colony of C57BL/6 isogenic strains of WT, CD19^-/-^, J_H_^-/-^, Aicda^-/-^, Rag1^-/-^, IgA^-/-^, and MHCII^-/-^ mice has been maintained by the Kubinak lab at the University of South Carolina. All animals used in the experiments described here are derived from this colony and were comprised of age-matched (8-week-old) male and female animals from each strain. A total of 30 mice were used for this study (n=6 per genotype and divided equally between males (n=15 total males used) and females (n=15 total females used)). Animals were reared and maintained under identical SPF conditions in a single room used to house the Kubinak mouse colony. All animals were maintained under constant environmental conditions (70°F, 50% relative humidity, 12:12 light:dark cycles) and were given *ad libitum* access to autoclaved drinking water and an irradiated soy-free mouse chow (Envigo; diet#2920X). All animal use strictly adhered to federal regulations and guidelines set forth by the University of South Carolina Institutional Animal Care and Use Committee (Approved Protocol#101580).

### Cell Isolations

(*Small bowel sample collection*) Animals were sacrificed and a 15cm section of small bowel (mid-jejunum to ileocecal valve) was collected in 15mL conical tubes containing 1X phosphate-buffered saline (PBS) and placed on ice. (*Tissue dissociation and IEC/ISC isolation*) The small bowel samples were placed on ice-cold 1X PBS-soaked paper towels, and Peyer’s patches were removed prior to downstream processing. Next, the tissue was flushed of luminal contents using 1X PBS and any connected mesenteric tissue and/or fat was removed. The cleaned tissue was washed by swirling in ice-cold 1X PBS, cut into small segments (~1cm), and then transferred to a sterile 50mL conical tube containing 15mL of cell dissociation buffer (1X HBSS with no Ca^++^ (GIBCO, 14175-095), 0.5M of EDTA (Invitrogen, 15575-038), 1 mM DTT (Thermofisher, J1539706), and 5% FBS (Corning, 35-011-CV). Segments were incubated at 37°C for 30 minutes with shaking at 200 revolutions per minute (rpm).After incubation, samples were briefly vortexed (5-10 seconds) and passed through a 100μm cell strainer into a new 50mL conical tube. The flow-through (containing IECs and ISCs) was centrifuged at 1500rpm for 10 minutes and the cell pellets were then resuspended in 500μL of complete RPMI media (RPMI 1640 supplemented with FBS, sodium pyruvate, non-essential amino acids, L-glutamine, penicillin-streptomycin, and β-ME). Cell counts were performed on isolated IECs. (*Enzymatic digestion and immune cell isolation*) The remaining tissue on the 100μm cell strainer was collected and transferred to a fresh 50mL conical tube containing 10mL of digestion buffer (1X HBSS with Ca^++,^ (GIBCO, 14025-092), 5% of FBS (Corning, 35-011-CV), 1 unit/mL Dispase (Worthington-biochem, 4942078001), 0.5mg/mL of Collagenase D (Roche, 11088882001), and 50μg/mL of DNase I (Worthington-biochem, LS002139). Samples were briefly vortexed to mix and were then incubated at 37°C for 20 minutes with shaking at 200rpm. Following incubation, samples were vortexed for 30-60 seconds (until no tissue was visible). Samples were then passed through a 40μM cell strainer into a fresh 50mL conical tube. A subsequent rinse with 5ml of ice-cold 1X PBS was swirled in the previous conical and passed through the strainer to collect and wash through any residual cells. Next, the samples were centrifuged at 2500rpm for 10 minutes, and the supernatant was discarded. The cell pellets were then resuspended in 40% Percoll (Cytiva, 17544501) and transferred to a fresh 15mL conical tube. Gently, 80% Percoll was pipetted underneath the 40% Percoll layer maintaining a distinct separation between layers. Subsequently, the samples were centrifuged at 2500rpm for 20 minutes without brake and acceleration. The samples were removed from the centrifuge and a pipette was used to remove the top 1mL of solution containing debris. Cells located at the 40%/80% interface were then transferred by pipette into a new 50mL conical tube. 50mL of ice cold 1X PBS was then added to the conical tube to wash cells. Samples were then centrifuged at 2500rpm for 10 minutes. Supernatant was discarded and cell pellets were resuspended in 500μL of complete RPMI. Cell counts were performed on isolated tissue-resident immune cells.

### Cell staining

*(Cell activation)* From each immune cell isolation, samples were divided into equal halves (250μL). One half was used to perform our 33-marker surface stain, and the other half was used to perform our 17-color T cell stain. Prior to staining for T cells, cells underwent activation. For the activation step, cells were suspended in complete RPMI media containing cell activation cocktail (Biolegend, 423303) and subsequently incubated at 37°C in a CO_2_ incubator for 4 hours. After incubation, cells were washed twice with column buffer (1X HBSS, 5mM EDTA, 5% FBS). To wash, cells were resuspended in 250uL of column buffer, mixed by gentle pipetting, and then spun at 1350rpm for 5 minutes. *(Fc-receptor blocking)* All immune cell stains incorporated an Fc-receptor block prior to staining. To do this, cells were incubated in 100μL of Fc-blocker reagent (Biolegend, 156603) for 10 minutes. (*Cell viability staining*) For live/dead cell discrimination, viability staining was performed using the Zombie-Aqua viability dye (live=ZombieAqua^-^; dead=ZombieAqua^+^). To do this, cells were washed with 1X PBS and subsequently resuspended in 100μL of 1X PBS containing the viability dye at a concentration of 1:500. Following a 15 minute incubation at room temperature, the cells were centrifuged at 1350rpm for 5 minutes and washed with column buffer (1X HBSS, 5% FBS, and 0.5 M EDTA). (*Surface markers staining*) Cells were resuspended in 100μL volumes of column buffer containing antibodies against relevant cell surface markers and incubated at room temperature in the dark for 20 minutes. After staining, cells were washed twice with column buffer (1X HBSS, 5mM EDTA, 5% FBS). To wash, cells were resuspended in 250uL of column buffer, mixed by gentle pipetting, and then spun at 1350rpm for 5 minutes. Cells used for the IEC/ISC panel and 33-marker immune panel were washed in staining buffer and then fixed in 500μl of a 1:1 ratio of column buffer and 4% paraformaldehyde (PFA). Fixed and stained cells were then kept in the dark at 4° C until analysis. Cells requiring intracellular staining (our T cell panel) were washed with permeabilization buffer (Biolegend, 421002) after surface staining and prior to fixation. (*Intracellular staining*) Cells were resuspended in 100μL volumes of intracellular fixation buffer (Biolegend, 420801) and incubated at RT for 20 minutes in the dark. Next, cells were centrifuged at 1350rpm for 5 minutes and the supernatant was discarded. Fixed cells were washed with permeabilization buffer twice, and then cells were resuspended in permeabilization buffer containing antibodies against relevant intracellular markers. After 30 minutes of room temperature incubation in the dark, cells were spun at 1350rpm for 5 minutes, supernatant was discarded, and pellets were washed twice with column buffer. After the final wash, cells were then fixed in 500μL of a 1:1 ratio of column buffer and 4% PFA. Fixed and stained cells were then kept in the dark at 4° C until ready to analyze. Supplementary Table T1 contains detailed information regarding the flow cytometry reagents and antibodies utilized in this study.

### Raw flow cytometry data handling

Flow cytometry analysis of prepared cells was conducted using a high-parameter BD FACSymphony A5 cell analyzer (IECs/ISCs) and a high-parameter spectrally-equipped BD FACSymphony A5SE cell analyzer (immune cells). Spectral compensation, t-distributed stochastic neighbor embedding (tSNE) data dimensionality reduction, and manual gating to define cellular subsets was performed on flow cytometry data using FlowJo Software (version 10.8.2, BD Biosciences). Briefly, a single compensated spectral spillover-spread-matrix was generated and applied to all sample data files. Manual gating to eliminate doublets and dead cells, and to identify major cellular lineages (erythroid (ter119^+^), endothelial (CD31^+^), epithelial (epithelial cell adhesion molecule (EpCAM^+^)), and immune (CD45^+^)) was performed and applied to all samples. For immune cell enumeration using our 33-marker general immunophenotyping panel, the liveCD45^+^ter119^-^CD31^-^EpCAM^-^ populations from all samples were concatenated into a single .fcs file for tSNE data dimensionality reduction. For immune cell enumeration using our 17-marker T cell panel, the liveCD45^+^CD3ε^+^NK1.1^-^ populations from all samples were concatenated into a single .fcs file for tSNE data dimensionality reduction. For IEC/ISC enumeration using our 10-marker IEC/ISC cell panel, the liveCD45^-^EpCAM^+^ populations from all samples were concatenated into a single .fcs file for tSNE data dimensionality reduction. Prior to dimensionality reduction of concatenated dataset, event-downsampling was performed to ensure equal contribution of events from each sample for tSNE construction. Manual gating was used to identify all cell subsets and were mapped onto constructed tSNE plots. An identical gating rubric was applied to all samples. Mouse models of primary antibody deficiency (IgA^-/-^, J_H_^-/-^, CD19^-/-^, Aicda^-/-^) and primary immunodeficiencies (MHCII^-/-^, Rag1^-/-^) served as important biological controls to validate gating strategies. Gating rubrics used to define each of the cellular subsets shown in this study are provided in Supplementary Figures S1-S6 and Supplementary Table T2.

### Small bowel enteropathy scoring

From each mouse, 1cm of the distal ileum was removed from mice during tissue collection from sacrificed animals. This tissue was cleaned as described above to remove luminal contents, mesenteric tissues, and fat, and was then subsequently placed in a 15mL conical tube containing 10% buffered-formalin for 48 hours. Tissue was submitted for processing to the USC Instrumentation Resource Facility Histology Core for embedding and H&E staining. Tissues were embedded in paraffin, sectioned, and H&E stained. Ileal sections were imaged with an EVOS microscope at 20X magnification. Areas of inflammation were scored blindly and based on three scoring metrics according to Erben et al.^17^: leukocytosis, mucosal architecture and overall extent of inflammation throughout the section. Multiple inflamed areas per section were selected for scoring based on clear orientation and quality of sectioning and staining. Leukocytosis was scored based on a combined metric of density (percent of cells per area of 4-5 well oriented villi and crypts) and severity (extent of expansion to various types of tissue, I.e. epithelial, mucosa, submucosa, muscularis scored 0-3). Mucosal architecture was scored based on a combined metric of villus to crypt ratios (scored 0-4) and structural damage (qualitatively given a score based on epithelial damage, abscesses, and irregularity shaped villi and crypts scored 0-4). A final disease score was calculated by adding the combined metrics described above (Leukocytosis + Mucosal architecture) and then multiplying by the extent of inflammation or the percentage of inflamed tissue throughout the entire 1cm section.

### Statistical analysis

Prism9.0 (GraphPad) was used for univariate pairwise statistical comparisons. Normality was assessed using the Shapiro-Wilk’s test. Means were used to indicate our measure of central tendency. For normally-distributed datasets involving comparisons between three or more groups, multiple hypothesis testing (with correction) was performed by applying a Dunnett test using the WT cohort as control (“all vs. WT” comparisons). For non-normally-distributed datasets, multiple hypothesis testing was performed by applying a Kruskal-Wallis test. All significant effects shown in figures with multiple comparisons reflect FDR-corrected p-values. For comparisons between two groups, a Student’s t-test was used for normally-distributed datasets and a Mann-Whitney U test was used for non-normally-distributed datasets. All multivariate statistics were performed using JMP16 Software (SAS). To minimize the effect of non-normally distributed datasets, all data analyses were performed on log_10_-transformed data. To obtain a numeric estimate of immune dysregulation (what we term as our “Immune Dysregulation Index (IDI)”) we calculated how dissimilar each antibody-deficient mouse strain is from the WT condition. We did this by comparing Mahalanobis distances generated by performing linear discriminant analysis (LDA) of all 55 cellular phenotypes using “mouse strain” as the response variable. Mahalanobis distances derived from independent LDAs were also used to determine the contribution of specific immune responses (e.g. innate responses, adaptive responses, IEC/ISC phenotypes, CD1d/MHCII expression, or effector T/iNKT cell responses) by incorporating subsets of relevant cellular phenotypes into respective LDA models (again using mouse strain (i.e. genotype) as our response variable). The Response Screening function in SAS performs multiple linear regressions between single ‘dependent x independent variable’ comparisons. For these analyses, the IDI was modelled as the dependent variable against independent cell phenotypes. False discovery rate (FDR) correction for multiple comparisons was applied to response screening analyses to account for multiple hypothesis testing. The Predictor Screening function in SAS is a method of bootstrap forest partitioning that involves recursive simultaneous modelling of all variables for identification of those that maximally explain variance in a response variable. This analysis was used to identify the relative contribution of each cellular phenotype in explaining variance in both immune dysregulation and disease severity. Multidimensional scaling (MDS) was used to visually represent relationships among individual animals based on cellular phenotypes and are provided as MDS plots. MDS plots were constructed in Microsoft Excel.

## Results

### Development of a cellular atlas of the murine small bowel using high-dimensional flow cytometry

From each mouse used in this study, we collected small bowels for histological analysis of enteropathy and to perform high-dimensional flow cytometry on isolated immune and epithelial cell subsets (**Fig 1A**). From each animal, independent isolations were conducted to collect cell fractions enriched in immune cells or epithelial cells. The average total cell counts obtained for downstream use from each isolation ranged between 2-4×10^6^ cells, with isolations from CD19^-/-^ and J_H_^-/-^ mice yielding more total cells than other strains (**Fig 1B**). On average, our cellular isolations yielded greater than 60% viable cells (**Fig 1C**). Sex had no effect on the total number or viability of cells obtained from our isolations (**Fig 1D**). To eliminate spectral interference of auto-fluorescent dead cells from our flow cytometry assay we incorporated the Zombie-Aqua viability dye into our flow panel and gated out dead/dying cells (Zombie-Aqua^+^) from downstream analyses (**Fig 1E**). To further enhance data quality, we also incorporate lineage markers in our initial parent gates in order to eliminate erythroid (ter119^+^) and endothelial (CD31^+^) cell contaminants from our analysis and to delineate between epithelial (EpCAM^+^), and immune (CD45^+^) cell subsets (**Fig 1E**). MDS analysis illustrates that genotype rather sex was the major driver of differences in the absolute abundance of major cell lineages (**Fig 1F**). While the absolute abundances of major cell lineages were generally consistent among WT, IgA^-/-^, and Aicda^-/-^ mouse strains, significant deviations were observed in CD19^-/-^ and J_H_^-/-^ mice. Specifically, CD19^-/-^ and J_H_^-/-^ mice had significantly higher numbers of immune, erythroid, and epithelial cells in their small bowels compared to WT mice (**Fig 1G**). The remaining cellular markers were used to develop a cellular atlas of immune/IEC/ISC phenotypes specific to the small bowel of mice that allow us to enumerate a total of 55 cell subsets (**Fig 1H**) (**Supplementary Table T2, Supplementary Fig S1-S6**).

**Fig 1.**
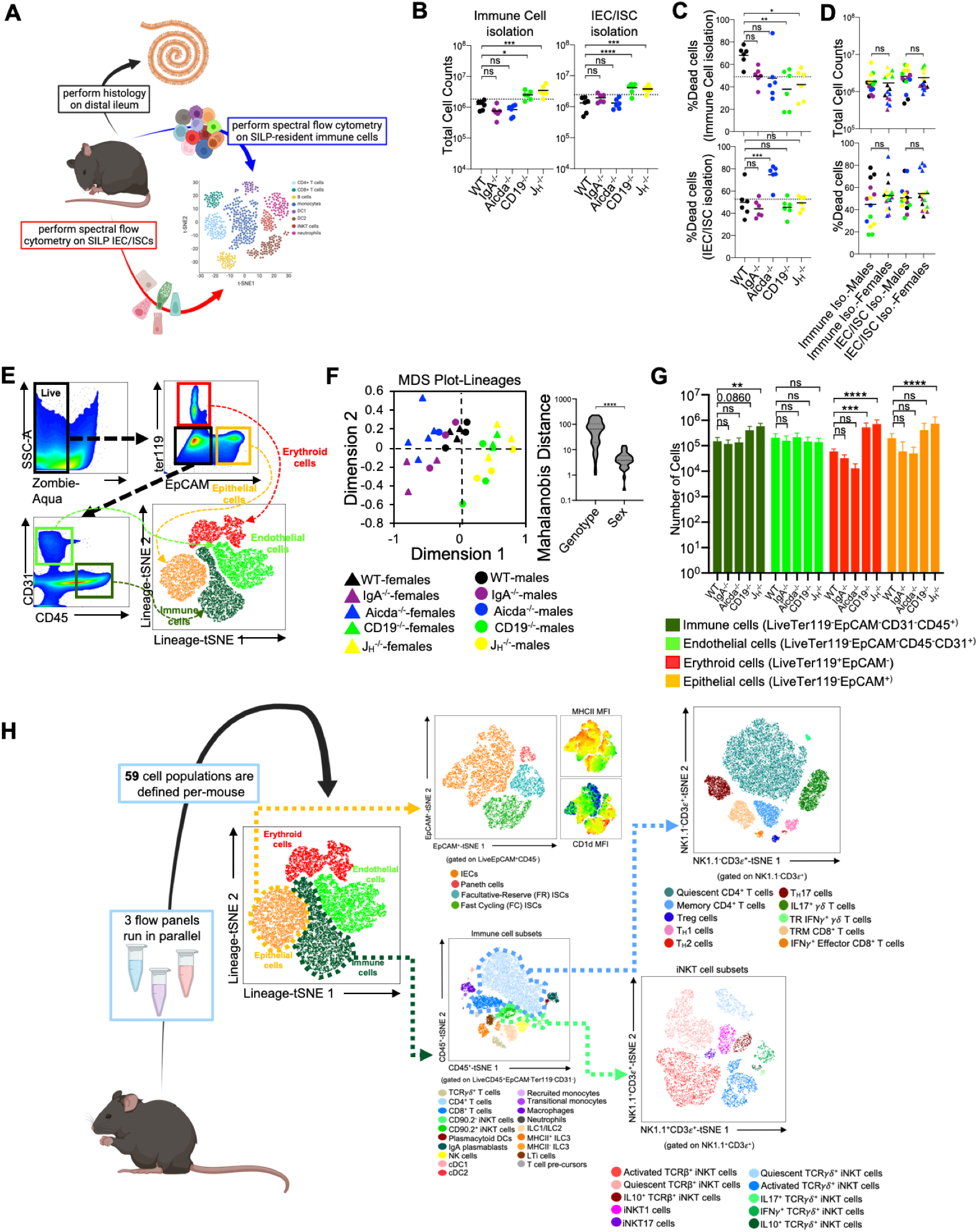
Development of a cellular atlas of the murine small bowel using high-parameter flow cytometry. **(A)** A schematic summarizing our experiment design is shown. **(B)** The total cell counts that are obtained using our two cell isolation strategies are shown. **(C)** Cell viability reflected as the percentage of dead cells is shown. **(D)** The distributions of total cell counts and cell viability are plotted by sex for each isolation strategy. **(E)** A representative gating strategy used to enumerate live cells and four major cell lineages is provided. **(F)** An MDS plot illustrating effect of genotype (marker colors) and sex (marker shapes) on the abundance of lineage^+^ cells is shown. **(G)** The absolute abundance of lineage^+^ cells are shown across genotypes. **(H)** An atlas of 59 cellular phenotypes that can be discriminated using our SI-customized high-parameter flow cytometry approach is provided. (B,C,G) Dunnett’s Test (“all vs. WT”), ns=p>0.05, *=p<0.05, **=p<0.01, ***=p<0.001, ****=p<0.0001. (D,F) Unpaired Student’s t-test, ns=p>0.05, ****=p<0.0001.

### PAD-induced immune dysregulation is associated with enhanced type I immunity and diminished iNKT cell responsiveness

The five mouse strains utilized in this study represent a spectrum of primary antibody deficiency (**Fig 2A**), which we predicted would result in a gradient of immune dysregulation. To test this prediction, we calculated the phenotypic divergence in immune responses (hereafter referred to as the immune dysregulation index (IDI)) between antibody-deficient and WT mice. The utility of this approach is three-fold. First, it allows us to generate a quantifiable estimate of immune dysregulation. Second, it allows us to identify which immunological and epithelial factors are primary drivers of this phenotype. Third, it allows us to identify immunological/epithelial factors that are predictive of disease across a range of underlying genetic predispositions. Based on this metric, we found that IDI scores generally agreed with our initial expectations. IgA^-/-^ mice representing a low IDI phenotype, Aicda^-/-^ mice an intermediate IDI phenotype, and J_H_^-/-^ and CD19^-/-^ high IDI phenotypes (**Fig 2B**). Our flow assay allows us to identify nineteen immune cell subsets (**Fig 2C**). Concatenation (pooling) of data derived from all animals allows us to visually depict shifts in the relative abundance of cells among mouse genotypes in the form of density plots (each plot represents an amalgam of events collected from six different mice of the same strain)(**Fig 2D**). The inclusion of knockout mouse strain density plots (RAG1^-/-^ and MHCII^-/-^ in this study) are provided to enhance rigor in the gating rubrics we used to define cell populations and were not included in any statistical analysis. As expected, significant differences were observed in the absolute abundance of multiple cell subsets across our antibody-deficient mouse strains (**Supplementary Fig S1, S2**). Collectively, multivariate analysis of these nineteen immune cell clusters demonstrates that immune dysregulation among antibody-deficient mouse strains is primarily driven by abnormal adaptive immune responses (**Fig 2E**). Specifically, the abundances of several T cell subsets (CD4^+^ T cells, T cell pre-cursors, and TCRγδ^+^ T cells) were positively correlated with IDI scores, whereas the abundances of most other cell subsets were not (**Fig 2F**). Thus, these latter differences are largely due to strain-dependent effects that are not shared drivers of immune dysregulation among mice.

**Fig 2.**
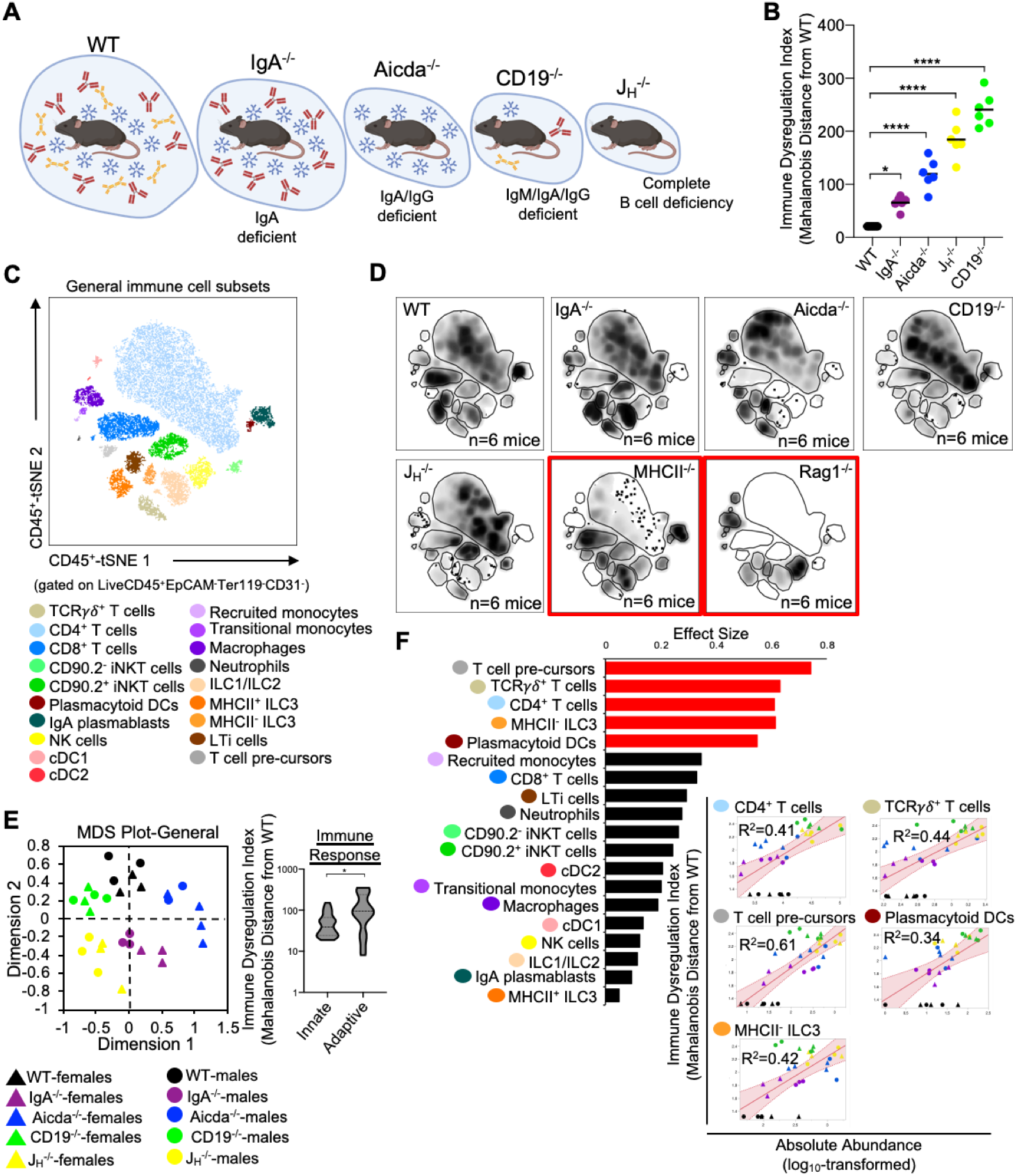
PAD-induced immune dysregulation is primarily associated with defects in T cell homeostasis in the small bowel. **(A)** PAD phenotypes and anticipated degree of immune dysregulation (size of circles) expected in mouse strains utilized in this study. **(B)** The degree of immune dysregulation (i.e. divergence in immune phenotype from the WT condition) among mouse strains is shown. Dunnett’s Test (“all vs. WT”), *=p<0.05, ****=p<0.0001. **(C)** A representative tSNE map of cellular subsets identified using our general immunophenotyping panel is shown. **(D)** Density tSNE plots highlighting differences in cellular distributions among genotypes are shown (each plot is generated from concatenated events from six mice per genotype). T and B cell deficient RAG1^-/-^ and CD4-deficient MHCII^-/-^ mice are provided as gating controls. **(E)** An MDS plot illustrating immunophenotypic variation among mouse strains (left), and the relative contribution of innate versus adaptive immune responses in driving immune dysregulation (right) are shown. Mann-Whitney U test, *=p<0.05. **(F)** The results of response screening analysis of immune phenotypes that significantly correlate (denoted by red bars) with the degree of immune dysregulation are shown. Significance reflects results of FDR-corrected p-values from multiple linear regression analysis of all 19 general immune cell subsets.

Previous studies from our lab support that primary antibody deficiency perturbs T cell homeostasis in the small bowel and that this may drive gut inflammation^18,19^. Results from our analysis above are consistent with this. To expand upon this observation, we designed our flow assay to include 17 cellular markers that allows us to discriminate T cell subsets. Ten conventional T cells (NK1.1^-^CD3ε^+^) subsets could be identified with this assay (**Fig 3A**). Overall, the relative abundance of cells was generally consistent among mouse strains with J_H_^-/-^ and CD19^-/-^ mice tending to have more T cells than other strains (**Fig 3B**). Notably, significant enrichment of effector T_H_17 cells was found to be a strain-dependent and CD19^-/-^-specific effect (**Supplementary Fig S3**), whereas enrichment of T_H_1 cells is a general observed feature in the small bowel of all antibody-deficient mouse strains (**Fig 3C**). The absolute abundance of all conventional T cell subsets was positively correlated with IDI scores, with T_H_1 responses having the largest effect (**Fig 3D**). Ten iNKT cell subsets (NK1.1^+^CD3ε^+^) subsets could also be identified with this assay (**Fig 3E, Supplementary Fig S4**). IDI scores were positively correlated with the abundance of several effector iNKT populations (iNKT1, iNKT17, and IL10^+^ iNKT) but were most strongly correlated with the abundance of quiescent (i.e. inactive) iNKT cell subsets (**Fig 3F, 3G**). Multivariate analysis of these twenty T/iNKT cell clusters demonstrates that immune dysregulation is more strongly driven by abnormalities in effector T cell responses rather than effector iNKT responses (**Fig 3H**).

**Fig 3.**
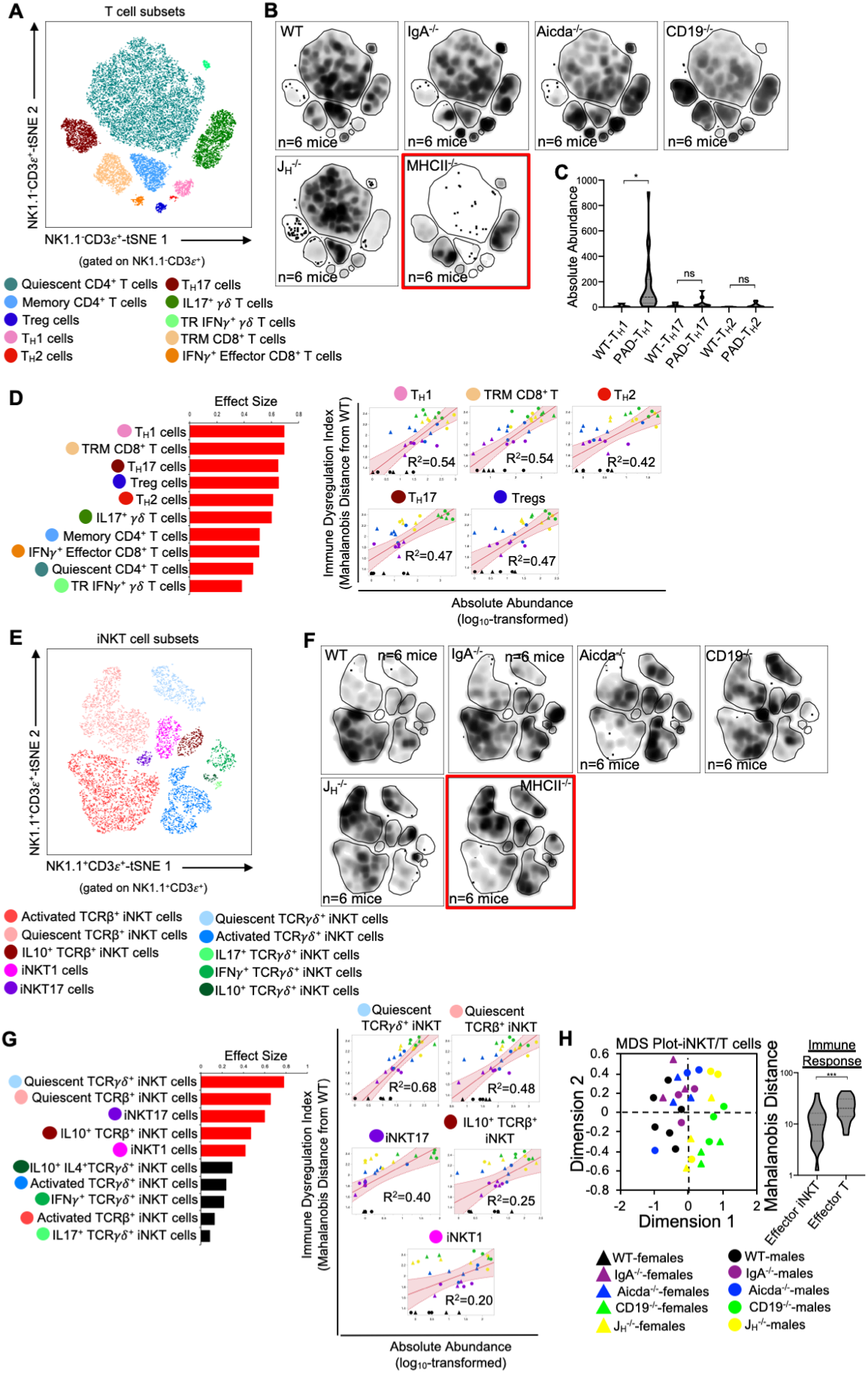
PAD-induced immune dysregulation is associated with enhanced type I immunity and iNKT cell quiescence. **(A)** A representative tSNE map of identifiable conventional T cell subsets is shown. **(B)** Density tSNE plots highlighting differences in absolute T cell subset abundances are shown (each plot is generated from concatenated events from six mice per genotype). CD4-deficient MHCII^-/-^ mice are provided only as gating controls. **(C)** Statistical comparisons of the absolute abundance of T_H_1, T_H_17, and T_H_2 cell between WT and PAD mice are shown. Kruskal-Wallis test, *=p<0.05, ns=p>0.05. **(D)** The results of response screening analysis of T cell phenotypes that significantly correlate (denoted by red bars) with the degree of immune dysregulation are shown. Significance reflects results of FDR-corrected p-values (FDR-corrected p-values <0.05) from multiple linear regression analysis of all 10 T cell subsets. The top 5 most significant correlations between T cell subset abundance and immune dysregulation are shown. **(E)** A representative tSNE map of identifiable iNKT cell subsets is shown. **(F)** Density tSNE plots highlighting differences in absolute iNKT cell subset abundances are shown (each plot is generated from concatenated events from six mice). CD4-deficient MHCII^-/-^ mice are provided only as gating controls. **(G)** The results of response screening analysis of iNKT cell phenotypes that significantly correlate (denoted by red bars) with the degree of immune dysregulation are shown. Significance reflects results of FDR-corrected p-values from multiple linear regression analysis of all 10 iNKT cell subsets. The top 5 most significant correlations between iNKT cell subset abundance and immune dysregulation are shown. **(H)** An MDS plot illustrating immunophenotypic variation among mouse strains based on T and iNKT cell phenotypes (left), and the relative contribution of effector T cell and effector iNKT cell responses in driving immune dysregulation (right) are shown. Mann-Whitney U test, ***=p<0.001.

### PAD-induced immune dysregulation is associated with aberrant antigen presentation by ISCs and IEC

We (and others) have previously shown that antibody-deficient mice have defects in bile acid biochemistry and lipid absorption in the small bowel^14,15,20^. While IECs and ISCs are known to present protein antigens via MHCII molecules and lipid antigens via CD1d molecules, it is currently unknown whether the balance between protein and lipid antigen presentation influences T cell homeostasis in the small bowel. To assess the relationship between immune cell homeostasis and epithelial cell antigen presentation we designed our flow assay to incorporate 10 cell markers that allowed us to identify MHCII- and CD1d-expressing Paneth cells, fast-cycling (FC) ISCs, facultative-reserve (FR) ISCs, and differentiated intestinal epithelial cells (IECs). A total of sixteen cellular phenotypes can be quantified using this set of markers (**Fig 4A, Supplementary Fig S5**). Overall, what we observed was a general deficiency in antibody-deficient mice in the number of IECs/ISCs expressing CD1d, and an enrichment in IECs/ISCs expressing MHCII (**Fig 4B, Supplementary Fig S6**). Multivariate analysis also revealed that shifts in the abundance of IECs/ISCs that expressed either MHCII or CD1d alone (single positive (SP)) were the strongest drivers of immune dysregulation compared to CD1d^-^MHCII^-^ double-negative (DN) or CD1d^+^MHCII^+^ double-positive (DP) IECs/ISCs (**Fig 4C, 4D**). Additionally, the abundance of CD1d^+^MHCII^-^ FC and FR ISCs was negatively correlated with immune dysregulation, which were the only two cellular phenotypes to do so (**Fig 4D**). Collectively, this data indicates that IECs and ISCs can assume one of four antigen-presenting phenotypes, that polarization towards MHCII-rather than or CD1d-mediated antigen presentation by IECs/ISCs is a feature of primary antibody deficiency in mice, and that ISCs focused on lipid antigen presentation may be central to maintaining immune homeostasis in the small bowel.

**Fig 4.**
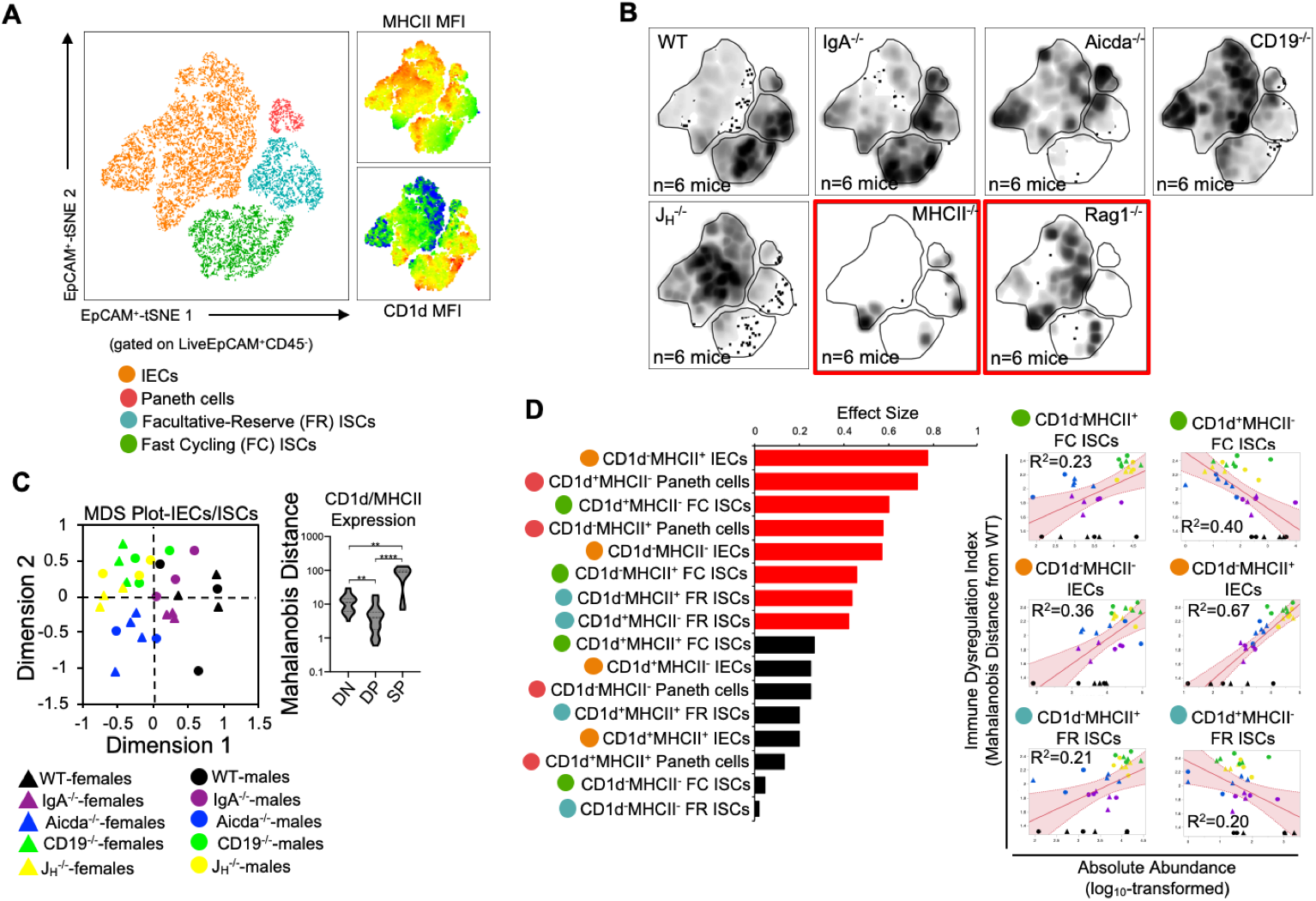
PAD-induced immune dysregulation is associated with aberrant antigen presentation by ISCs and IECs.. **(A)** A representative tSNE map of identifiable ISC and IEC subsets is shown. Four lineages (IEC, Paneth cells, FC ISCs, and FR ISCs) were subsequently gated based on their expression of MHCII and CD1d antigen presenting molecules. Heatmaps of MHCII and CD1d expression are provided to orient the reader to major shifts in the abundance of antigen-presenting IECs and ISCs shown in 4B. **(B)** Density tSNE plots highlighting differences in absolute IEC/ISC subset abundances are shown (each plot is generated from concatenated events from six mice). T and B cell deficient RAG1^-/-^ and CD4-deficient MHCII^-/-^ mice are provided as gating controls. **(C)** An MDS plot illustrating immunophenotypic variation among mouse strains based on IEC and ISC antigen-presenting phenotypes (left), and the relative contribution of differences in the absolute abundances of double-negative (CD1d^-^MHCII^-^)(<u>DN</u>), double-positive (CD1d^+^MHCII^+^)(<u>DP</u>), and single-positive (CD1d^+^MHCII^-^ or CD1d^-^MHCII^+^)(SP) cell subsets in driving immune dysregulation (right) are shown. Kruskal-Wallis test, **=p<0.01, ****=p<0.0001. **(D)** The results of response screening analysis of immune phenotypes that significantly correlate (denoted by red bars) with the degree of immune dysregulation are shown. Significance reflects results of FDR-corrected p-values from multiple linear regression analysis of all 16 IEC/ISC subsets analyzed.

### Diminished lipid antigen presentation by ISCs may allow pathologic type 1 immune responses to develop in the small bowels of antibody-deficient mice

Mice from all four antibody-deficient strains develop small bowel enteropathy (**Fig 5A**) with the extent of disease similar between male and female animals (**Fig 5B**). Among genotypes, a gradient of disease severity was observed which was that significantly correlated with IDI scores (**Fig 5C**). Multivariate analysis of all 55 cell subsets indicated that disruption to adaptive immunity was again the most important driver of overall immune dysregulation in mice, followed by defects in IEC/ISC phenotypes, and finally innate immune responses (**Fig 5D**). Additionally, when all 55 cell subsets were simultaneously compared for their ability to predict IDI and disease severity, we found that ten cell subsets were highly predictive of both (**Fig 5E**). Collectively, these subsets reflected type 1 immune responses (e.g. the abundance of T_H_1 cells, IFNγ^+^ CD8^+^ T cells, plasmacytoid DCs), iNKT cell quiescence, and CD1d-mediated lipid antigen presentation (**Fig 5E**). As noted above, shifts in the relative abundance of IECs/ISCs expressing either CD1d or MHCII (single-positive (SP)) were major discriminating phenotypes among mouse strains (**Fig 4C**). While the relative abundance of DN, DP, and SP cells were generally consistent among the four IEC/ISC cell subsets analyzed, the absolute abundance of CD1d SP cells differed. Specifically, most of the CD1d SP cells are found in the FC ISC subset of WT mice (**Fig 5F, 5G**). In contrast, antibody-deficient mice (particularly Aicda^-/-^, CD19^-/-^ and J_H_^-/-^) were deficient in these cells and instead enriched for MHCII SP cells in both their FC ISC and IEC subsets (**Fig 5G**). Finally, we analyzed the correlation between the abundance of ISCs/IECs and T cell/iNKT cell subsets identified as being predictive of disease in 5E. This analysis revealed that the abundance of CD1d-expressing ISCs (and especially FC ISCs) is inversely correlated with type 1 immune responses and iNKT cell quiescence (**Fig 5H**). This data suggests that under normal conditions, polarization towards lipid antigen presentation in ISCs favors iNKT cell activation that suppresses type 1 immunity in the SI.

**Fig 5.**
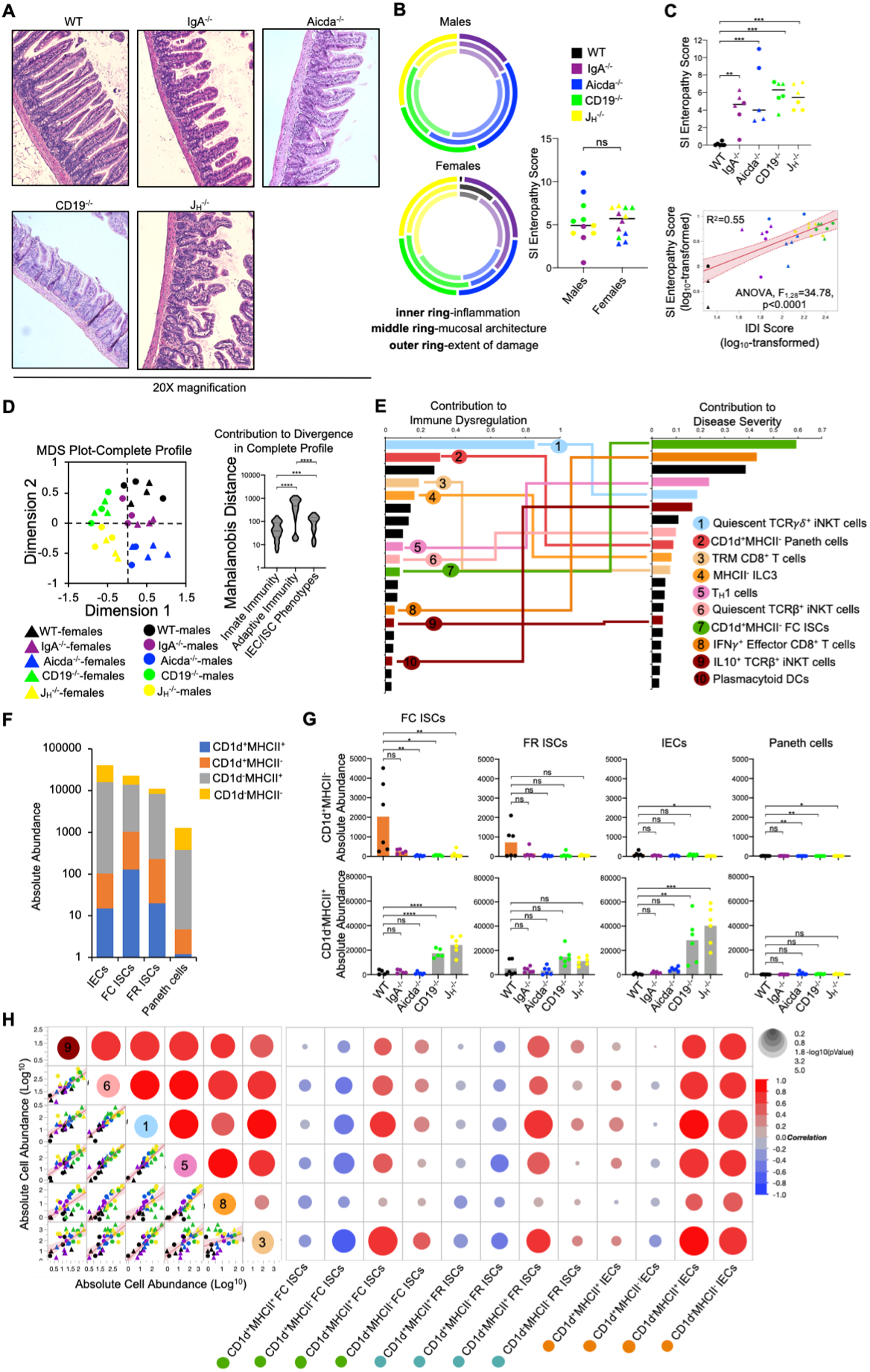
Diminished lipid antigen presentation by ISCs may allow pathologic type 1 immune responses to develop in the small bowels of antibody-deficient mice. **(A)** Representative H&E images (20X magnification) of distal ileal sections from PAD mouse models are shown. (B) Radial plots are provided to show similarity between genders in each disease parameter. Cumulative enteropathy scores are compared between males and females. Student’s t-test, ns=p>0.05. **(C)** The severity of small bowel enteropathy among mouse strains is shown (upper plot). Dunnett’s Test (“all vs. WT”), **=p<0.01, ***=p<0.001. A regression analysis of relationship between immune dysregulation and disease severity is shown. ANOVA. **(D)** An MDS plot reflecting divergence among individuals based on their cumulative immunophenotypes is shown (left) and the relative effects of cumulative innate immune responses, adaptive immune responses, or IEC/ISC antigen presentation in driving immune dysregulation are compared (right). Kruskal-Wallis test, ***=p<0.001, ****=p<0.0001. **(E)** Results of predictive screening analysis of immune phenotypes that best predict IDI and disease severity scores among mouse strains are shown. The top 20 significant discriminators for each outcome (IDI or disease) are plotted with overlapping cellular phenotypes highlighted by connecting lines. “Contribution” reflects the contribution of each variable to variance among mouse strains in IDI or disease severity. **(F)** Stacked barplots showing the absolute abundance of antigen presenting phenotypes in IEC/ISC subsets are shown. **(G)** The absolute abundance of single positive cells (CD1d^+^MHCII^-^ or CD1d^-^MHCII^+^) among the four IEC/ISC subsets is shown. Dunnett’s Test or Tukey’s test (“all vs. WT”), ns=p>0.05, *=p<0.05, **=p<0.01, ***=p<0.001, ****=p<0.0001. **(H)** A correlation matrix summarizing results of multiple linear regression analysis of relationships between IEC/ISC antigen presentation phenotypes and T cell/iNKT cell subsets identified in (E) is shown.

## Discussion

The laboratory mouse is an unparalleled tool for studying inborn errors of immunity as it is a tractable model with the most comprehensive set of reagents developed to study immune system function. Given the confounding variables associated with cohort studies in humans and the oversimplification of *in vitro* models, the laboratory mouse is the only experimental system that can adequately capture the dynamic and multifaceted nature of vertebrate immunity and the disease consequences of dysregulated inflammation. Here, we demonstrate a novel application of flow cytometry, facilitated by recent advancements in this technology, that may yield novel insights into underlying mechanisms of disease.

Immune dysregulation is the hallmark feature of primary immunodeficiency but lacks a generalizable definition because of the heterogenous nature of this spectrum of immunological disorders. A quantifiable metric of immune dysregulation based on a comprehensive set of functional readouts of an individuals’ immune response would be beneficial for several reasons. It would allow us to identify what immune responses are dysregulated, quantify the magnitude of dysregulation, and use these values to determine the value of each immune biomarker as a predictor of disease risk/severity. Recent advances in flow cytometry now provide a rapid and cost-effective approach to acquire composite readouts of immune function in a tissue-specific manner. This can be utilized (in tandem with other classical diagnostic tests) to acquire a more complete understanding of a patients immunophenotype.

In our study, we explicitly defined immune dysregulation as deviation in the overall immunophenotype of an individual from baseline. WT mice served as our baseline condition and Mahalanobis distance was used to estimate deviation in the immune responses of individual mice. Based on this approach, we show that immune dysregulation in the small bowel covaries with the expected degree of immunodeficiency across a series of mouse strains and is positively correlated with a physiological outcome (small bowel enteropathy). Our approach also reveals that defects in lipid antigen presentation by ISCs may destabilize homeostatic T cell responses and skew towards inflammation, which highlights a potentially novel axis of inflammatory disease unique to the small bowel. Currently, there are over 300 clinically recognized primary immunodeficiencies that have been associated with numerous genetic mutations^1,4^. However, variability in penetrance and pleiotropic effects of these varied mutations diminish the value of such diagnostics in the identification of potentially generalizable therapeutic approaches. From this perspective, methods like ours may be more fruitful as they facilitate rapid screening of genetically-disparate individuals to identify shared functional responses that affect disease outcomes.

Chronic non-infectious enteropathies are commonly observed in patients with several different primary antibody deficiencies. For instance, chronic diarrhea is a common gastrointestinal manifestation that is frequently observed in patients with X-linked agammaglobulinemia (XLA), common variable immunodeficiency (CVID), selective IgA deficiency (SIgAD), and IgG subclass deficiency^21,22^. Additionally, in mice, we and others have shown that several different genetic models of antibody deficiency develop inflammatory disease in the gut, which may involve both the small bowel^14,15,20,23^ and colon^24,25^. These animal studies have demonstrated that antibody-deficient mice share transcriptional, metabolic, and histological features observed in antibody-deficient patients^19,20,26-28^. While the underlying mechanisms driving disease in antibody-deficient patients are currently undefined, the manifestations of immune dysregulation have been extensively studied in CVID patients. Studies restricted to peripheral T and B cell-mediated responses in CVID patients demonstrated that patients present with fewer naïve CD4^+^ and CD8^+^ T cell in their blood ^29-31^, and more activated cells^32^. However, these T cell phenotypes were not observed in less clinically-severe forms of primary antibody deficiency including IgG-subclass deficiency and selective IgA deficiency^32^, consistent with the degree of immune dysregulation being less severe in such patients. Collectively, results from several CVID cohort studies suggest that defects in homeostatic T cell responses triggered by aberrant molecular crosstalk between T cells and antigen presenting cells may be primary driver of gastrointestinal inflammation^28,33^.

IECs and ISCs serve unconventional roles as APCs that can coordinate T cell responses in the gut^34,35^. Several pieces of evidence support that regulation of inflammatory T_H_1 responses through IEC/ISC MHCII-mediated antigen presentation may be relevant to the pathogenesis of enteropathy in antibody-deficient mice and humans. Several studies using *in vitro* modelling or mouse models have shown that IFNγ can induce MHCII expression by IECs, and conversely that MHCII expression by IECs can promote intestinal T_H_1-driven inflammation (reviewed in^34^). However, one study has suggested that MHCII expression by IECs and ISCs regulates the behavior of antigen-experienced T cells rather than serve to initially activate naïve T cells^36^. Previous work in mice has shown that MHCII-mediated antigen presentation by IECs/ISCs can regulate T_H_1 immune responses, though whether it positively or negatively regulates this response appears to be highly context-dependent^37,38^. Conversely, T_H_1 responses have also recently been shown to regulate ISC renewal, differentiation, and pool size^39^. For example, it was shown that ISC MHCII expression resulted in accumulation and activation of crypt-resident CD4^+^ T cells. Moreover, inflammatory CD4^+^ T cells (T_H_1, T_H_2, T_H_17) and their canonical cytokines (IFNγ, IL-13, or IL17) were shown to limit ISC renewal capacity and consequently pool size, while regulatory T cell abundance was shown to play an opposing role. In our study, we find that severe antibody deficiency is associated with enhanced abundance of MHCII-expressing IECs and elevated T_H_1 responses. Two studies have shown that T_H_1 responses are exacerbated in CVID patients. In a cohort study of 29 CVID patients presenting with non-infectious disease complications, plasma protein profiling demonstrated that plasma type 1 cytokine levels (IL-12/IFNγ) were associated with more severe disease outcomes which included enteropathy^40^. Moreover, in an independent study of 13 CVID patients, observed increases in lamina propria lymphocytosis in CVID patients presenting with enteropathy was linked to increased production of type 1 cytokine (IL-12/IFNγ) synthesis by T cells isolated from these patients and compared to asymptomatic controls^41^. Whether MHCII-mediated antigen presentation by IECs/ISCs is perturbed in antibody-deficient humans and drives pathologic T_H_1 responses is unknown.

Despite their low contribution to overall immunophenotypic variation among antibody-deficient mice, iNKT cell responses appear to be an important driver of immune dysregulation and disease severity in our model. Specifically, we found a positive correlation between the number of quiescent iNKT cells and immune dysregulation, disease severity, and T_H_1 cells in antibody-deficient mice. In both mice and humans, iNKT cells are CD1-restricted T cells whose activation are regulated by lipid antigens^42^. While mice have a single CD1 molecule (CD1d), humans have four (CD1a-d)^43,44^. iNKT cells contribute to homeostatic immunity in the gut, and play an important role in regulating type 1 immune responses through their rapid production of immunoregulatory cytokines (discussed in depth by Crosby & Kronenberg^45^). In WT mice, we found that most CD1d-expressing cells were found within the FC ISC subset. Additionally, and in contrast to patterns of MHCII expression on IECs, results from our experiments indicate that antibody deficiency is associated with reduced CD1d expression in FC ISCs. Given the role iNKT cells play in homeostatic barrier defense, it is possible that defective CD1d-mediated ISC-iNKT cell interactions lead to iNKT cell quiescence and a resulting compensatory T_H_1 response. Disruption to normal lipid metabolism/absorption by gut ISCs could explain this effect. CVID patients have known defects in lipid homeostasis (recently reviewed in^46^), and in two specific studies of CVID patients (that also included the use of antibody-deficient mouse models), bulk RNA sequencing of small intestinal tissues revealed a signature of lipid malabsorption uniquely associated with CVID patients presenting with small bowel enteropathy^14,15^. Similar effects were observed in antibody-deficient mouse strains used in these studies. Two subsequent studies by our group using CD19^-/-^ mice reported a similar observation and linked this with defects in bile acid homeostasis in the small bowel^19,20^. Bile acid composition in the small bowel regulates the absorption of dietary lipids and lipid-soluble vitamins (A and D), and a recent study has linked CVID with bile acid malabsorption (though the study was limited in terms of participants)^47^. Collectively, these observations suggest that defects in lipid metabolism/absorption could destabilize the iNKT-T_H_1 cell axis in the gut to drive enteropathy.

Several factors contribute to difficulties in studying gut mucosal immune responses in humans, and especially within the small bowel. First, obtaining samples from patients poses a significant challenge, leading researchers to rely mainly on inferences drawn from the cellular makeup of blood samples. Second, previous studies of the cellular basis of inflammatory disease in antibody-deficient patients have relied on flow cytometry panels based on a handful of markers which by necessity require major assumptions regarding which cell types are worth investigating and whether/how well inferences drawn from blood peripheral blood phenotyping reflects tissue-specific inflammatory responses. Our results highlight the clear benefit of high-parameter flow cytometry for maximizing the amount of information that can be obtained from small tissue samples. Additionally, the high-parameter flow cytometry panel we have developed and described in this study represents the most comprehensive flow cytometry assay for assessing small-bowel-specific cellular heterogeneity in mice. The immediate value of the development of such an assay is three-fold. First, we can simultaneously acquire physiological readouts (histological disease assessment in this study) as well as data on 55 cellular phenotypes from the SI of every mouse, which drastically reduces the number of animals needed for experimentation; a key ethical consideration in the design of animal studies. Second, our ability to generate information on all of these parameters on a per-mouse basis affords the ability to perform multivariate analyses to quantify effect sizes of key variables (e.g. sex, genotype, cell type, immune response) on disease severity, and to quantify the strength of linear relationships among all continuous variables. Third, high-parameter flow cytometry is a cost-effective means of quickly assessing diverse cellular responses, which aids in the refinement of experiments involving more costly genomics technologies like single cell RNA sequencing.

## Data Availability

Supplementary Data File 1 provides an excel sheet containing all of the raw data used in this analysis.

## Author contributions

J.L.K. designed and oversaw all experiments, performed data analysis, and wrote the first and final drafts of this manuscript. J.L.K., A.D.M., and R.A.W.B. equally contributed to the development of the flow cytometry assay described in this study. A.D.M., and R.A.W.B. performed all experiments, analyzed data, and co-wrote the first and final drafts of this manuscript. M.N. and P.N. supported this project through a COBRE (P20GM103641) pilot award, provided access to a BD FACSymphonsy A5SE, and assisted in manuscript revision.

## Figure captions

**Fig 1. Development of a cellular atlas of the murine small bowel using high-parameter flow cytometry. (A)** A schematic summarizing our experiment design is shown. **(B)** The total cell counts that are obtained using our two cell isolation strategies are shown. **(C)** Cell viability reflected as the percentage of dead cells is shown. **(D)** The distributions of total cell count and cell viability are plotted by sex for each isolation strategy. **(E)** A representative gating strategy used to enumerate live cells and four major cell lineages is provided. **(F)** An MDS plot illustrating effect of genotype (marker colors) and sex (marker shapes) on the abundance of lineage^+^ cells is shown. **(G)** The absolute abundance of lineage^+^ cells are shown across genotypes. **(H)** An atlas of 59 cellular phenotypes that can be discriminated using our small-bowel-customized high-parameter flow cytometry approach is provided. (B,C,G) Dunnett’s Test (“all vs. WT”), ns=p>0.05, *=p<0.05, **=p<0.01, ***=p<0.001, ****=p<0.0001. (D,F) Unpaired Student’s t-test, ns=p>0.05, ****=p<0.0001.

**Fig 2. PAD-induced immune dysregulation is primarily associated with defects in T cell homeostasis in the small bowel. (A)** PAD phenotypes and anticipated degree of immune dysregulation (size of circles) expected in mouse strains utilized in this study. **(B)** The degree of immune dysregulation (i.e. divergence in immune phenotype from the WT condition) among mouse strains is shown. Dunnett’s Test (“all vs. WT”), *=p<0.05, ****=p<0.0001. **(C)** A representative tSNE map of cellular subsets identified using our general immunophenotyping panel is shown. **(D)** Density tSNE plots highlighting differences in cellular distributions among genotypes are shown (each plot is generated from concatenated events from six mice per genotype). T and B cell deficient RAG1^-/-^ and CD4-deficient MHCII^-/-^ mice are provided as gating controls. **(E)** An MDS plot illustrating immunophenotypic variation among mouse strains (left), and the relative contribution of innate versus adaptive immune responses in driving immune dysregulation (right) are shown. Mann-Whitney U test, *=p<0.05. **(F)** The results of response screening analysis of immune phenotypes that significantly correlate (denoted by red bars) with the degree of immune dysregulation are shown. Significance reflects results of FDR-corrected p-values from multiple linear regression analysis of all 19 general immune cell subsets.

**Fig 3. PAD-induced immune dysregulation is associated with enhanced type I immunity and diminished iNKT cell responsiveness. (A)** A representative tSNE map of identifiable conventional T cell subsets is shown. **(B)** Density tSNE plots highlighting differences in absolute T cell subset abundances are shown (each plot is generated from concatenated events from six mice per genotype). CD4-deficient MHCII^-/-^ mice are provided only as gating controls. **(C)** The results of response screening analysis of T cell phenotypes that significantly correlate (denoted by red bars) with the degree of immune dysregulation are shown. Significance reflects results of FDR-corrected p-values from multiple linear regression analysis of all 10 T cell subsets. **(D)** Statistical comparisons of the absolute abundance of T_H_1, T_H_17, and T_H_2 cell between WT and PAD mice are shown. Kruskal-Wallis test, *=p<0.05, ns=p>0.05. **(E)** A representative tSNE map of identifiable iNKT cell subsets is shown. **(F)** Density tSNE plots highlighting differences in absolute iNKT cell subset abundances are shown (each plot is generated from concatenated events from six mice). CD4-deficient MHCII^-/-^ mice are provided only as gating controls. **(G)** The results of response screening analysis of iNKT cell phenotypes that significantly correlate (denoted by red bars) with the degree of immune dysregulation are shown. Significance reflects results of FDR-corrected p-values from multiple linear regression analysis of all 10 iNKT cell subsets. **(H)** An MDS plot illustrating immunophenotypic variation among mouse strains based on T and iNKT cell phenotypes (left), and the relative contribution of effector T cell and effector iNKT cell responses in driving immune dysregulation (right) are shown. Mann-Whitney U test, ***=p<0.001. **(I)** Significant correlations (FDR-corrected p-values <0.05) between T cell and iNKT cell subsets with immune dysregulation are shown.

**Fig 4. PAD-induced immune dysregulation is associated with aberrant antigen presentation by ISCs and IECs. (A)** A representative tSNE map of identifiable ISC and IEC subsets is shown. Four lineages (IEC, Paneth cells, FC ISCs, and FR ISCs) were subsequently gated based on their expression of MHCII and CD1d antigen presenting molecules. Heatmaps of MHCII and CD1d expression are provided to orient the reader to major shifts in the abundance of antigen-presenting IECs and ISCs shown in 4B. **(B)** Density tSNE plots highlighting differences in absolute IEC/ISC subset abundances are shown (each plot is generated from concatenated events from six mice). T and B cell deficient RAG1^-/-^ and CD4-deficient MHCII^-/-^ mice are provided as gating controls. **(C)** An MDS plot illustrating immunophenotypic variation among mouse strains based on IEC and ISC antigen-presenting phenotypes (left), and the relative contribution of differences in the absolute abundances of double-negative (CD1d^-^MHCII^-^)(<u>DN</u>), double-positive (CD1d^+^MHCII^+^)(<u>DP</u>), and single-positive (CD1d^+^MHCII^-^ or CD1d^-^ MHCII^+^)(SP) cell subsets in driving immune dysregulation (right) are shown. Kruskal-Wallis test, **=p<0.01, ****=p<0.0001. **(D)** The results of response screening analysis of immune phenotypes that significantly correlate (denoted by red bars) with the degree of immune dysregulation are shown. Significance reflects results of FDR-corrected p-values from multiple linear regression analysis of all 16 IEC/ISC subsets analyzed.

**Fig 5. PAD-driven SI enteropathy is associated with defects in CD1d-mediated regulation of homeostatic iNKT cell responses in the small intestine. (A)** Representative H&E images (20X magnification) of distal ileal sections from PAD mouse models are shown. (B) Radial plots are provided to show similarity between genders in each disease parameter. Cumulative enteropathy scores are compared between males and females. Student’s t-test, ns=p>0.05. **(C)** The severity of small bowel enteropathy among mouse strains is shown (upper plot). Dunnett’s Test (“all vs. WT”), **=p<0.01, ***=p<0.001. A regression analysis of relationship between immune dysregulation and disease severity is shown. ANOVA. **(D)** An MDS plot reflecting divergence among individuals based on their cumulative immunophenotypes is shown (left) and the relative effects of cumulative innate immune responses, adaptive immune responses, or IEC/ISC antigen presentation in driving immune dysregulation are compared (right). Kruskal-Wallis test, ***=p<0.001, ****=p<0.0001. **(E)** Results of predictive screening analysis of immune phenotypes that best predict IDI and disease severity scores among mouse strains are shown. The top 20 significant discriminators for each outcome (IDI or disease) are plotted with overlapping cellular phenotypes highlighted by connecting lines. “Contribution” reflects the contribution of each variable to variance among mouse strains in IDI or disease severity. **(F)** Stacked barplots showing the absolute abundance of antigen presenting phenotypes in IEC/ISC subsets are shown. **(G)** The absolute abundance of single positive cells (CD1d^+^MHCII^-^ or CD1d^-^MHCII^+^) among the four IEC/ISC subsets is shown. Dunnett’s Test or Tukey’s test (“all vs. WT”), ns=p>0.05, *=p<0.05, **=p<0.01, ***=p<0.001, ****=p<0.0001. **(H)** A correlation matrix summarizing results of multiple linear regression analysis of relationships between IEC/ISC antigen presentation phenotypes and T cell/iNKT cell subsets identified in (E) is shown.

