## Supplementary material for "Studying the cellular basis of small bowel enteropathy using high-parameter flow cytometry in mouse models of primary antibody deficiency": A Supplemental Figures

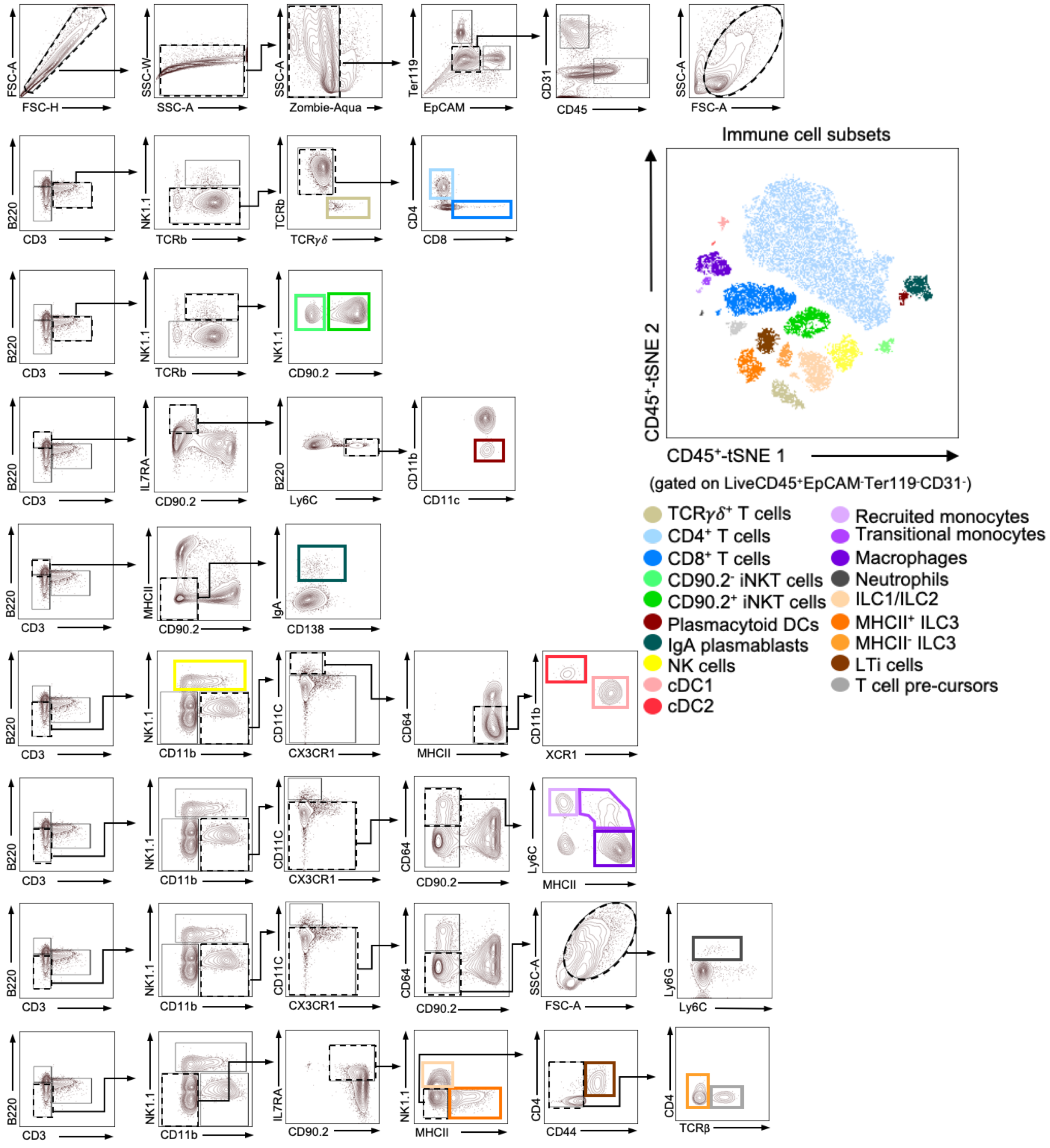


**Supplementary Figure S1. Gating rubric for immune cells.** Gating rubrics used to calculate absolute abundance of immune cell subsets are shown. Terminal gates defining tSNE cell clusters are color-coded. tSNE plot used in Figure 2A and population names corresponding to terminal gates are provided.

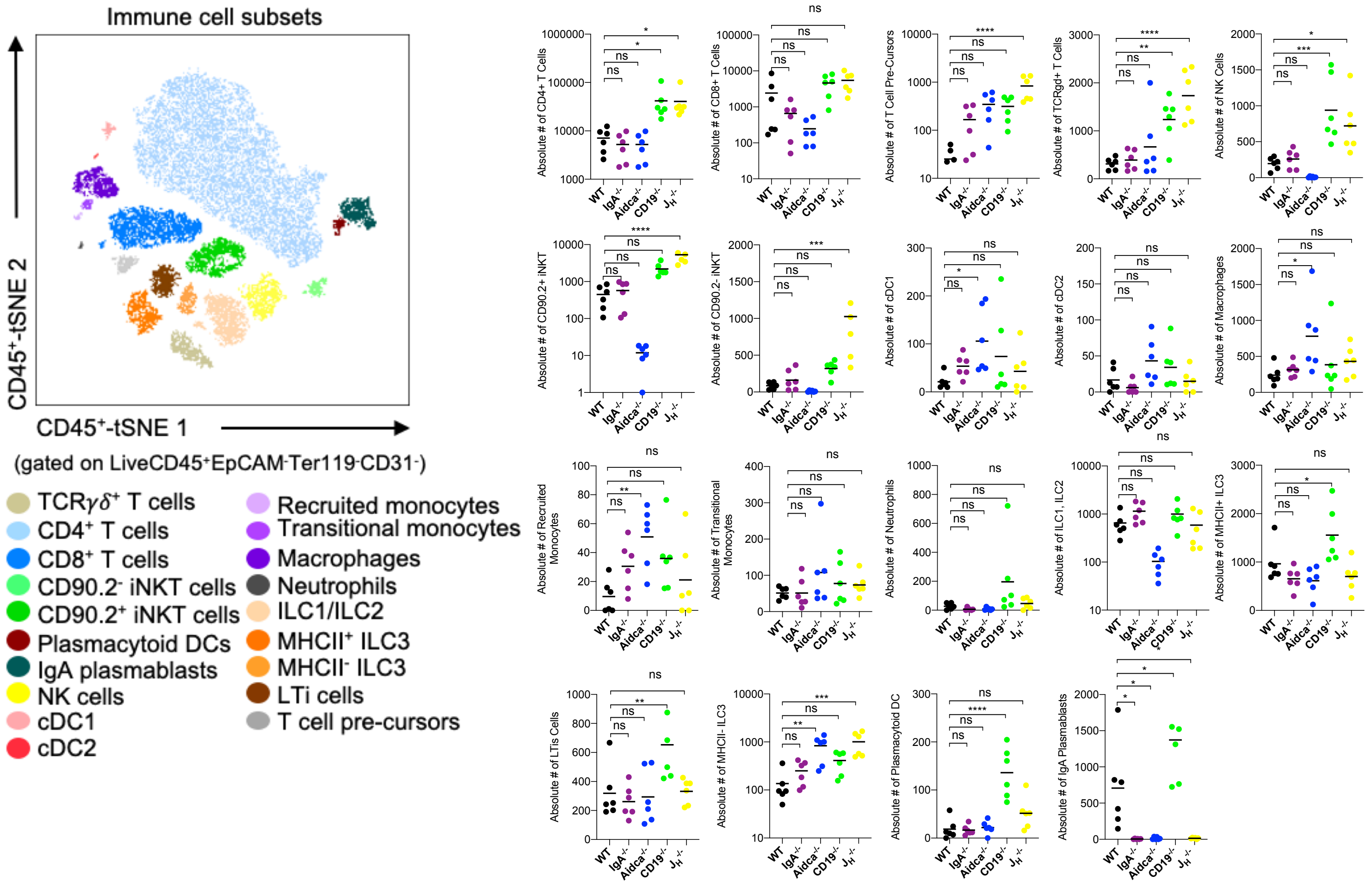


**Supplementary Figure S2. Statistical comparisons of all cellular subsets derived from our general immune panel.** Dunnett's Test ("all vs. WT"), ns=p>0.05, *=p<0.05, **=p<0.01, ***=p<0.001, ****=p<0.0001.

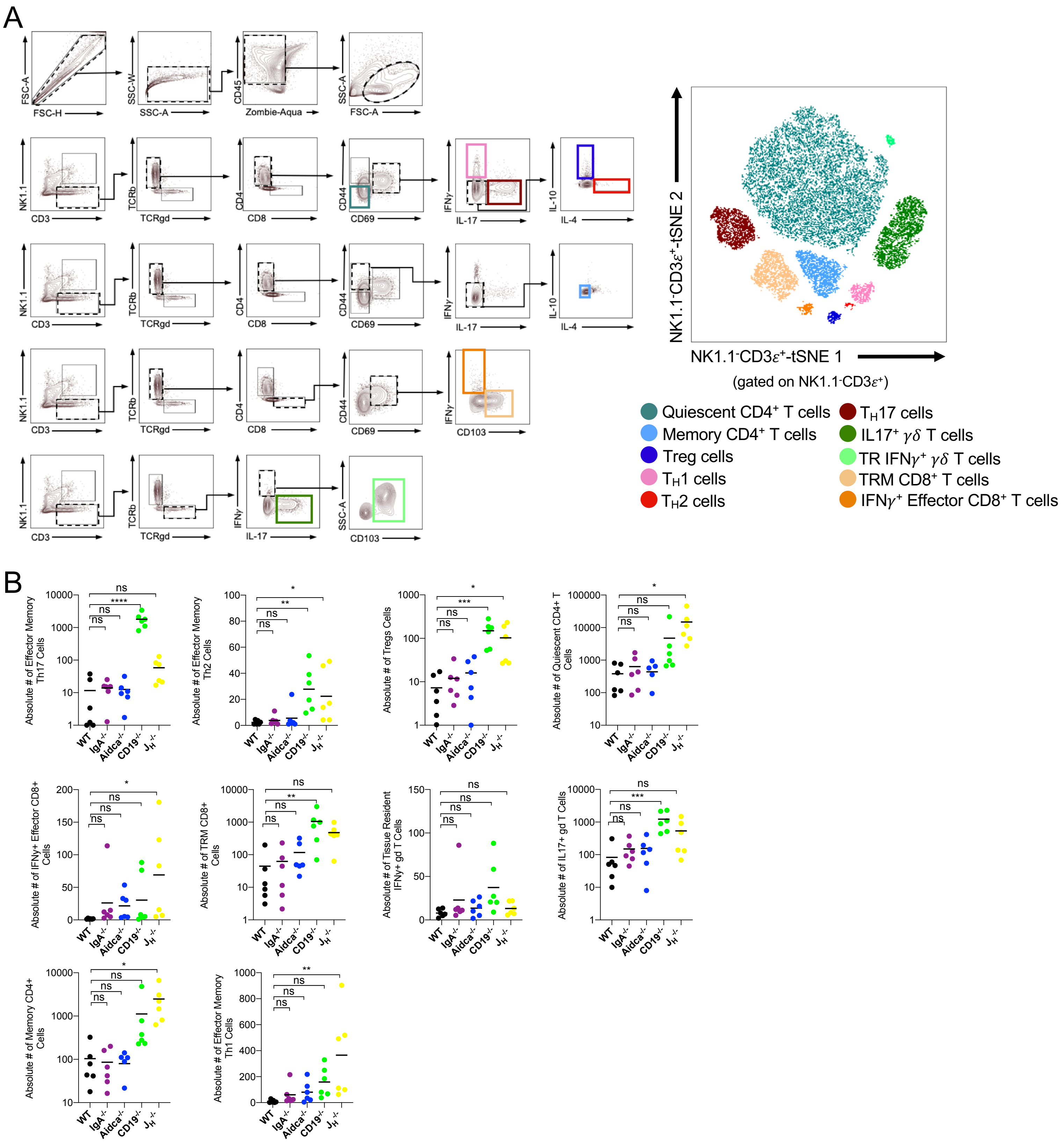


**Supplementary Figure S3 Gating rubric and statistical comparisons of conventional T cells. (A)** Gating rubrics used to calculate absolute abundance of conventional T cell subsets are shown. Terminal gates defining tSNE cell clusters are color-coded. tSNE plot used in Figure 3A and population names corresponding to terminal gates are provided. **(B)** Statistical comparisons of conventional T cell subsets are provided. Dunnett's Test ("all vs. WT"), ns=p>0.05, *=p<0.05, **=p<0.01, ***=p<0.001, ****=p<0.0001.

A

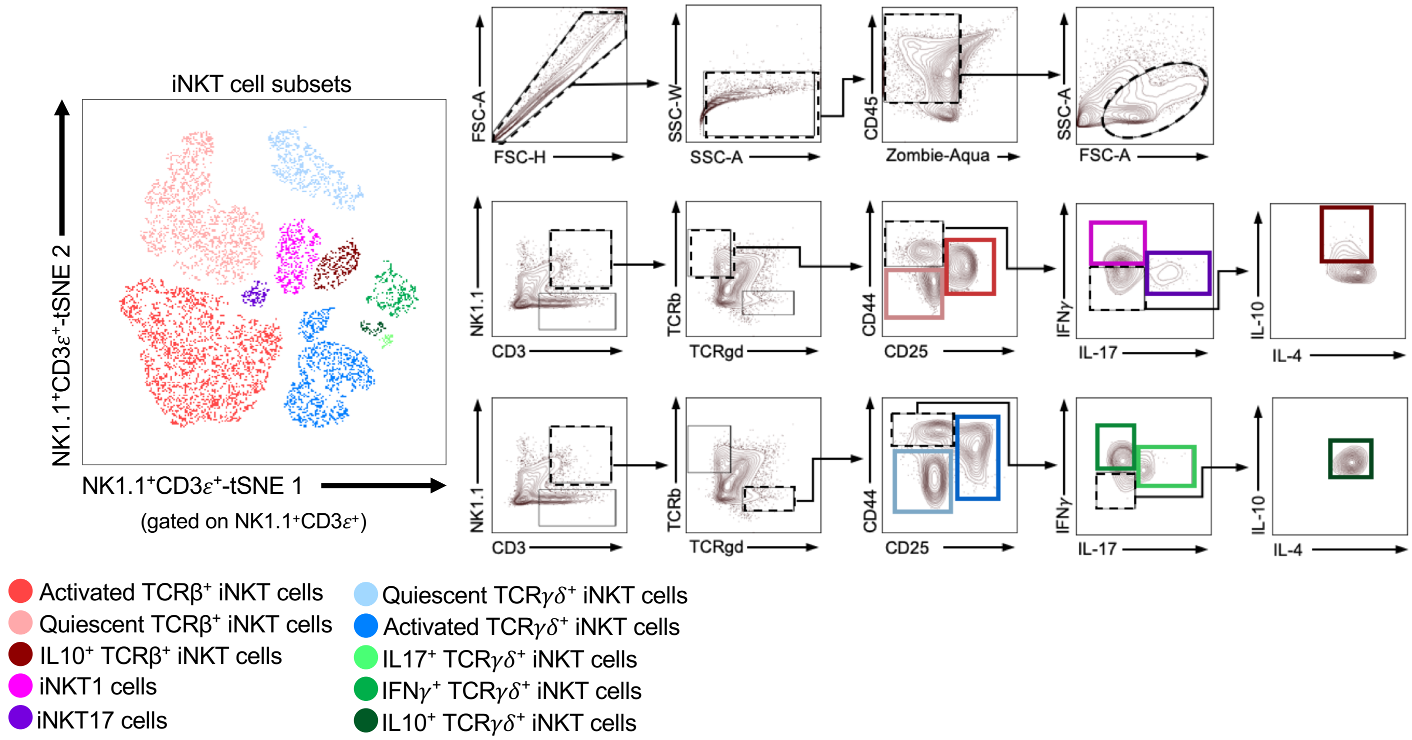


B

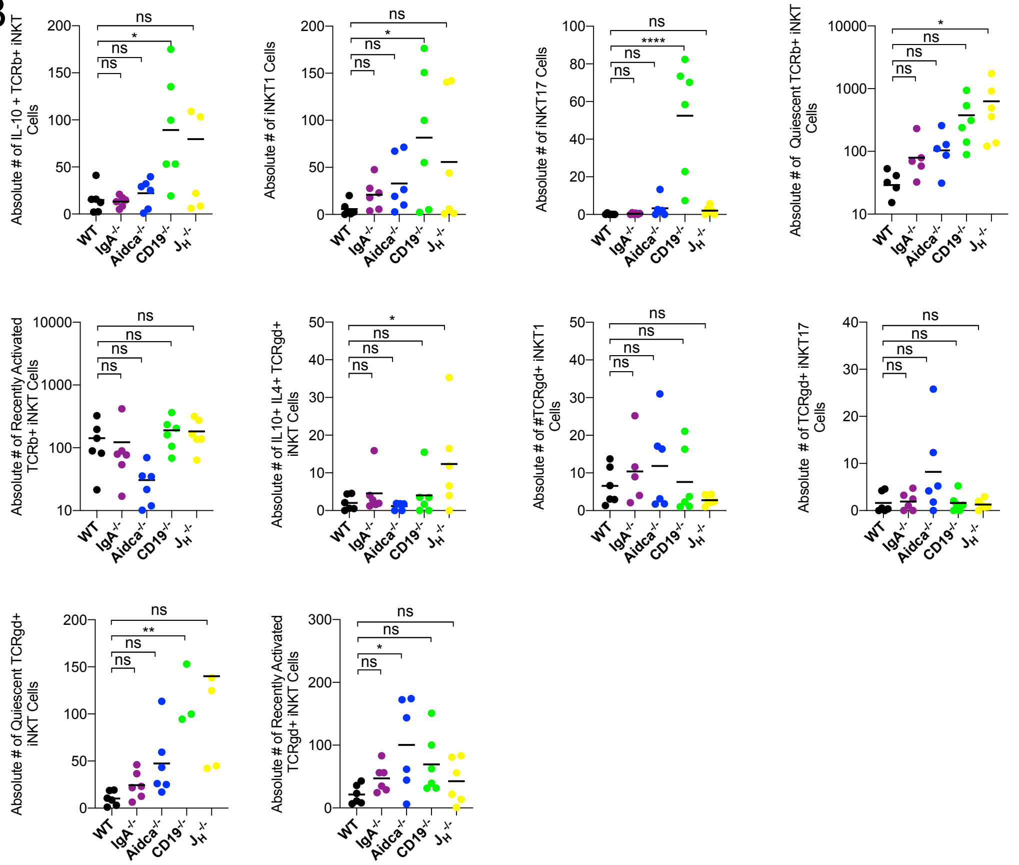


**Supplementary Figure S4. Gating rubric and statistical comparisons of invariant NKT cells. (A)** Gating rubrics used to calculate absolute abundance of invariant NKT cell subsets are shown. Terminal gates defining tSNE cell clusters are color-coded. tSNE plot used in Figure 3A and population names corresponding to terminal gates are provided. **(B)** Statistical comparisons of invariant NKT cell subsets are provided. Dunnett's Test ("all vs. WT"), ns=p>0.05, *=p<0.05, **=p<0.01, ***=p<0.001, ****=p<0.0001.

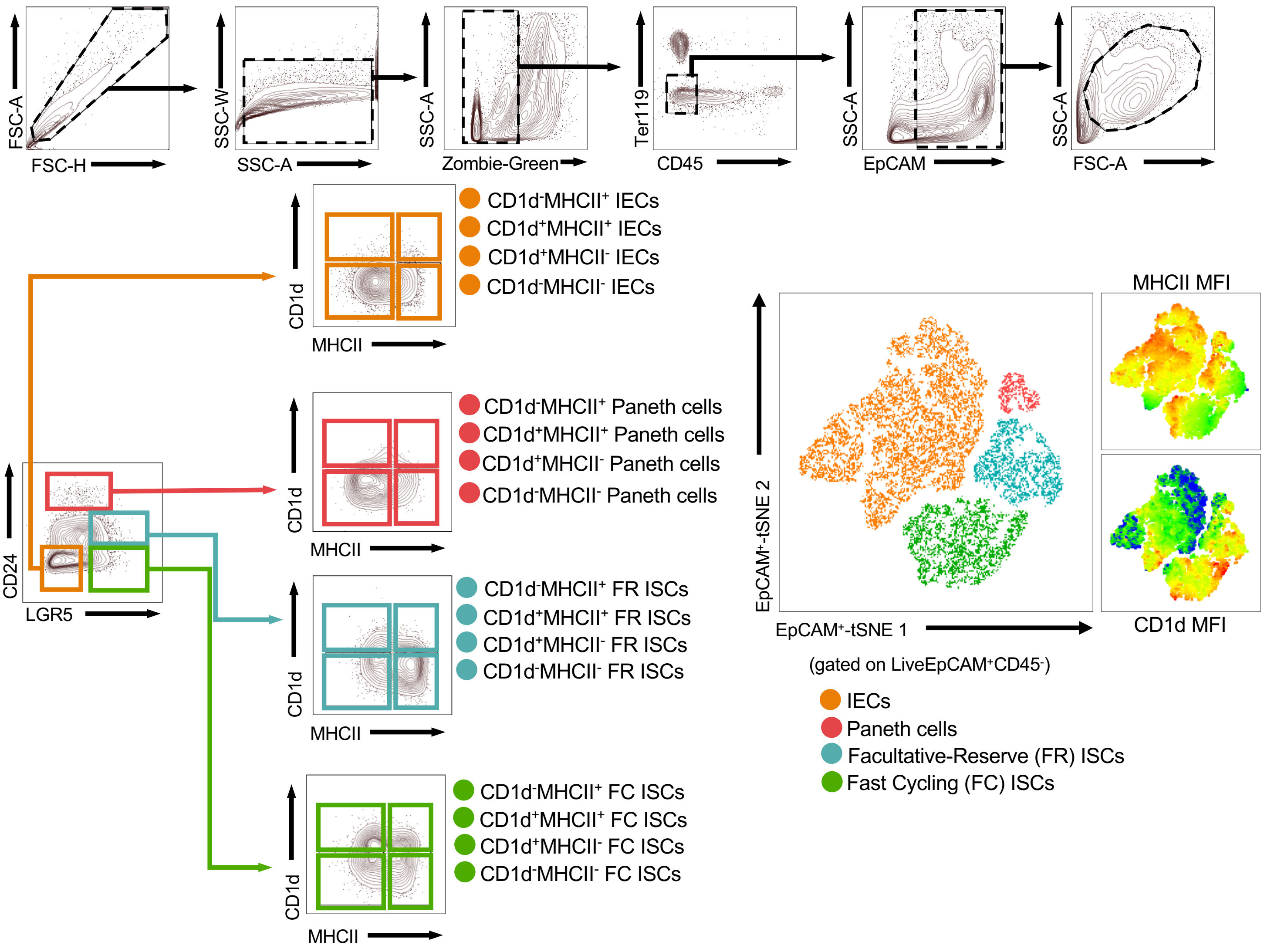


**Supplementary Figure S5 Gating rubric for IECs and ISCs.** Gating rubrics used to calculate absolute abundance of IECs and ISCs are shown. Terminal gates defining tSNE cell clusters are color-coded. tSNE plot used in Figure 4A and population names corresponding to terminal gates are provided.

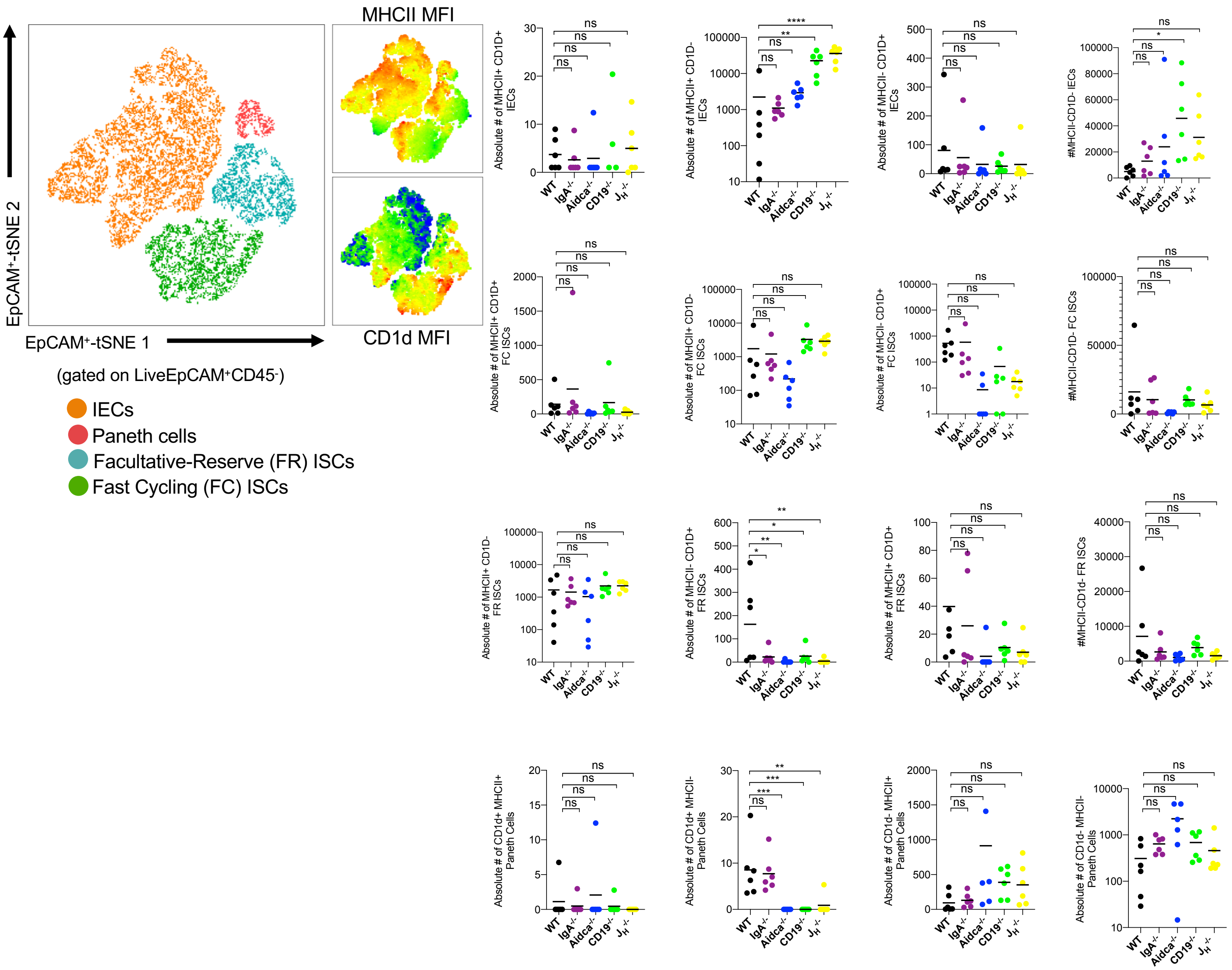


**Supplementary Figure S6. Statistical comparisons of all cellular subsets derived from our IEC/ISC panel.** Dunnett's Test ("all vs. WT"), ns=p>0.05, *=p<0.05, **=p<0.01, ***=p<0.001, ****=p<0.0001.
